# Go west: population genomics reveals unexpected population fluctuations and little gene flow in Western Hemisphere populations of the predatory lady beetle, *Hippodamia convergens*

**DOI:** 10.1101/2023.05.11.540381

**Authors:** Arun Sethuraman, Schyler O. Nunziata, Angela Jones, John Obrycki, David W. Weisrock

**Author notes:** Current address: United States Department of Agriculture (Schyler O. Nunziata). Current address: Duke University (Angela Jones).

## Abstract

The convergent lady beetle, *Hippodamia convergens*, is used extensively in augmentative biological control of aphids, thrips, and whiteflies across its native range in North America, and was introduced into South America in the 1950s. Overwintering *H. convergens* populations from its native western range in the United States are commercially collected and released across its current range in the eastern U.S., with little knowledge of the effectiveness of augmentative biological control using *H. convergens*. Here we use a novel ddRADseq-based SNP/haplotype discovery approach to estimate its range-wide population diversity, differentiation, and recent evolutionary history. Our results indicate (1) significant population differentiation among eastern U.S., western U.S., and South American populations of *H. convergens*, with (2) little to no detectable recent admixture between them, despite repeated population augmentation, and (3) continued recent population size expansion across its range. These results contradict previous findings using microsatellite markers. In light of these new findings, the implications for the effectiveness of augmentative biological control using *H. convergens* are discussed. Additionally, because quantifying the non-target effects of augmentative biological control is a difficult problem in migratory beetles, our results could serve as a foundation cornerstone in improving and predicting the efficacy of future releases of *H. convergens* across its range.

## Introduction

Genetic data are important sources of information for gaining insight into historical and recent evolutionary history of species that have had their populations established or augmented by large-scale releases of individuals (Kajita et al., 2012, Li et al., 2019, 2020, 2021, Sethuraman et al., 2020, Sentis et al., 2022, Jones et al., 2023). Population genomics, in particular, provides a powerful approach towards understanding how genetic variation has been impacted on a recent time scale (Nunziata et al., 2017; Maigret et al., 2020), including the time frame encompassing human-mediated movements and augmentations of populations. These insights can be important for at least two primary reasons. First, the pathway to successful pest suppression for many biocontrol introductions is often unknown. Second, by characterizing the degree to which population structure has evolved across the native range of a species over long-term time scales, and by estimating the impact of gene flow and population admixture between native and translocated populations (e.g., Li et al., 2020), the context for interpreting the non-target impacts of introduction efforts becomes available. For example, loss of genetic variation through biocontrol efforts could limit the potential for populations to adapt to future conditions, ultimately decreasing long-term fitness of released populations (Rhymer and Simberloff 1996; Roderick and Navajas 2003; Frankham 2005). Alternately, human-mediated releases may act to homogenize populations via admixture, and thereby reduce the potentially deleterious effects of localized pervasive inbreeding thus reinforcing the need to track the effects of gene flow and admixture due to human-mediated releases of species.

Large-scale releases of translocated or captive-raised organisms have been commonly used worldwide throughout the last century, including for conservation purposes (e.g., reintroduction or relief of inbreeding depression), commercial enterprises (e.g., logging, aquarium pet trade), sporting activities (e.g., hunting and fishing), and for biological control (Laikre et al., 2010). Despite the pervasiveness of these mass releases, non-target effects of translocations of native species are largely ignored in policy and research. While population augmentations can be demographically and economically beneficial, they can also adversely impact intraspecific genetic variation by causing the breakdown of local adaptations, loss of genetic variability, and the reshaping of population structure (Terui et al., 2023, Laikre et al., 2010, Simberloff and Stilling 1996, Wajnberg et al., 2001).

There is a long history of using insect predators for biological control, with billions of individuals being released over the past 100 years (Van Lenteren et al., 2003). Natural enemies are released to control pest populations, with or without the goal of permanent establishment of populations (Hajek and Eilenberg 2018, Heimpel and Mills 2017). Predatory coccinellids are widely used in biological control, with numerous native and exotic lady beetle species sold commercially and distributed throughout North America (Obrycki and Kring 1998). Population genetic studies have been useful in reconstructing population history of introduced lady beetle species that have successfully established populations in their introduced range (Lombaert et al., 2010; Kajita et al., 2012; Sethuraman et al., 2017). Less studied, but equally important is characterizing the population dynamics of a native species of lady beetle used in biological control (but see Sethuraman et al., 2015, 2017).

Convergent lady beetles, *Hippodamia convergens,* are one of the most abundant and widespread *Hippodamia* species in North and Central America, ranging from southern Canada and throughout the US, and into central America (Gordon 1985, Global Biodiversity Information Facility [www.GBIF.org], Lost Lady Beetle website [www.lostladybug.org]). This species has been established in its non-native range in South America, including in Peru and Chile, where millions of individuals were introduced over multiple years (Hagen et al., 1976, Global Biodiversity Information Facility [www.GBIF.org]). First used as biological control agents in the early 1900’s (Carnes, 1912), *H. convergens* is currently one of the most commonly used natural predators for control of aphids and whiteflies in the U.S. (van Lenteren 2003). Millions of adult individuals are collected by commercial insectaries annually from overwintering sites in California and sold for release (Dreistadt & Flint 1996; Obrycki and Kring 1998) throughout the U.S. A recent study employing seven microsatellite loci found California genotypes admixed into eastern U.S. populations with weak population structure across the range, a potential non-target effect of augmentation (Sethuraman et al., 2015). Alternatively, this pattern could be the result of natural movements. *Hippodamia convergens* is capable of long-distance migrations (Rankin and Rankin, 1980), often dispersing multiple miles (Hagan 1962).

In this study, we expand upon previous research by using genome-scale data sampled across a larger geographic range of *H. convergens*. Using large numbers of single-nucleotide polymorphisms (SNPs), we can refine our understanding of population structure and the genetic effects of releasing large numbers of western U.S. individuals across the larger range of the species. Further, SNP-based demographic inference allows for the testing of complex evolutionary histories, including timing and origins of population introductions, asymmetric levels of migration, and population size history (McCoy et al., 2014; Fraser et al., 2015).

Here, we used genome-wide SNP markers to examine genetic diversity and population genetic structure of *H. convergens* across its native range in the United States, and an introduced population in Chile. The aims of this study were to (1) examine the population structure of *H. convergens* across its range, (2) compare levels of genetic diversity, and (3) examine the population history including human-mediated and historic gene flow, population size history, and timing of population founding. Based on previous studies and history of population augmentation, we hypothesized: (1) admixture of western U.S. genotypes into eastern U.S. populations, (2) increases in contemporary gene flow, consistent with augmentation, and (3) a founder effect with corresponding low genetic diversity of the Chilean population.

## Methods

### Sampling, ddRAD Sequencing, and SNP genotyping

We collected genome-wide SNP data from 83 *H. convergens* individuals sampled from 16 sites broadly distributed throughout their known US range and 8 individuals sampled from one site in Chile representing a putative recently established population in South America (Figure 1; Table S1). This also included 18 individuals from two commercial suppliers in Arizona and California (Arbico and Rincon-Vitova, respectively). Locality information was not available for these purchased individuals; however, we assume they were collected from geographical locations in Arizona and California and represent native populations from these areas. For each sampling location we collected 1 to 12 individuals; however, some sampling locations were geographically close and pooled for analyses (Table S1).

**Figure 1:**
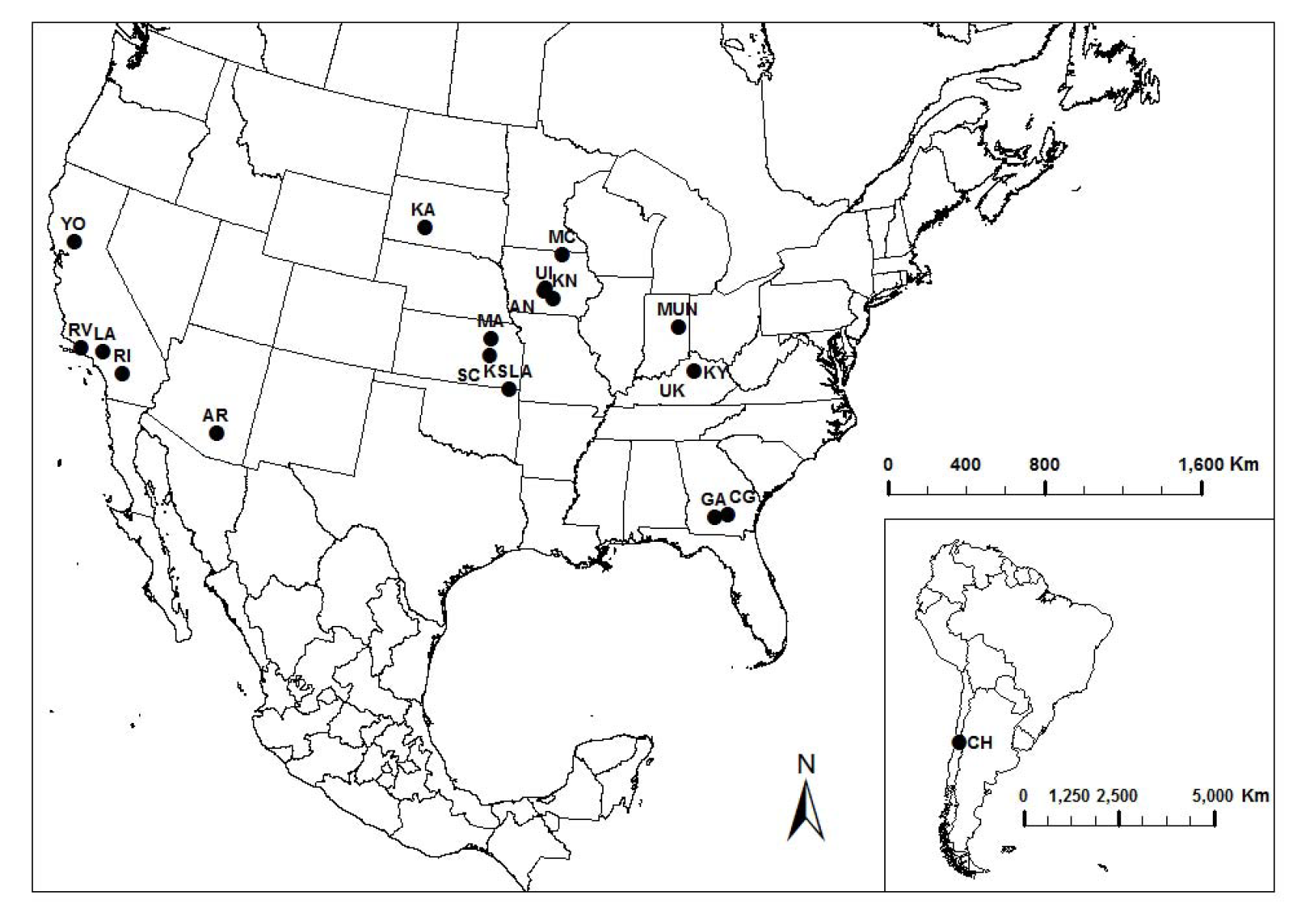
Geographical map of *Hippodamia convergens* sampling locations utilized in this study.

We performed double digest restriction-site associated DNA (ddRAD) sequencing using the protocol outlined in Peterson et al., 2012. Briefly, we extracted DNA from whole individuals (after gut removal) using a Qiagen DNEasy tissue kit (Qiagen, Valencia, CA, USA). We digested 1000 ng of DNA with the restriction enzymes *Eco*RI and *Nla*III. Custom inline barcodes and P1/P2 adapters were ligated onto the digested DNA fragments and uniquely barcoded samples were pooled and size-selected between 473 and 579 bp using a Pippin Prep (Sage Science). Size-selected DNA was then amplified using High-Fidelity DNA Polymerase (Bio-Rad) with 12 cycles. During PCR, a unique Illumina index sequence and Illumina sequencing primers were incorporated on ends of samples, so that each individual had a unique combination of inline and index sequences. All samples were then pooled and sequenced on a single lane of Illumina HiSeq 2500 at Florida State University’s College of Medicine using 150 bp paired-end reads. To increase sequence diversity in the initial portions of sequence reads, the library was sequenced using an additional 1% spike in of PhiX control library.

We used standard Illumina chastity filtering of our sequence data and performed downstream processing using STACKS v2.0Beta6 (Catchen et al., 2011, 2013). The *process_radtags* script was used to perform demultiplexing and quality-filtering. The *denovo_map.pl* script was used with default settings to identify unique RAD loci and call SNPs. The *populations* script was used to generate population datasets with loci that occurred in at least half of the sampling sites, in at least 50% of individuals at each site (r = 0.5), and with a minimum coverage of 10 reads (m = 10). After an initial run of the STACKS pipeline we excluded 8 individuals due to excessive missing data (> 50% missing genotypes across loci; Table 1S Supporting Information), bringing the total number of populations sampled down to 17. Further filtering was performed using vcftools v0.1.16 (Danecek et al., 2011) to retain bi-allelic SNPs genotyped across at least 50% of all individuals. Data was filtered to exclude SNPs with minor allele frequency (MAF) less than 0.05, which have been shown to create biases in quantifying genetic connectivity (Roesti et al., 2012; Benestan et al., 2015). To avoid the inclusion of linked sites, only a single SNP was sampled from a locus. Sequence (haplotypic) data were also generated for analyses of evolutionary history (see description below). Additional de novo assembly and filtering based on more conservative filters (PHRED Q score of ≥ 33, maximum of 5 low quality bases per read, minor allele frequency of ≥ 0.05, maximum of 8 indels, minimum of 4 samples per locus, maximum missing data of 25% per locus) using iPyRAD (Eaton et al., 2020) was also performed to obtain a smaller subset for population structure analyses.

**Table 1.** Summary statistics as calculated in the *populations* script in STACKS with 17 populations and 9824 SNPs. N = average number of individuals genotyped at each locus, private = number of variable sites unique to each population, H_obs_ = observed homozygosity.

| Region | Symbol | N | Private | $H_{obs}$ | $\pi$ | $F_{IS}$ |
| --- | --- | --- | --- | --- | --- | --- |
| West | AR | 2.54 | 183 | 0.960 | 0.066 | 0.043 |
| West | LA | 7.77 | 599 | 0.960 | 0.071 | 0.100 |
| West | RI | 3.20 | 253 | 0.962 | 0.069 | 0.058 |
| West | RV | 9.56 | 769 | 0.959 | 0.070 | 0.103 |
| West | YO | 4.70 | 363 | 0.960 | 0.072 | 0.074 |
| East | AN | 2.62 | 284 | 0.956 | 0.073 | 0.049 |
| East | CG | 4.89 | 589 | 0.957 | 0.078 | 0.084 |
| East | GA | 1.00 | 123 | 0.957 | 0.043 | 0.000 |
| East | KA | 1.62 | 189 | 0.955 | 0.058 | 0.021 |
| East | KN | 1.64 | 220 | 0.957 | 0.057 | 0.020 |
| East | KSLA | 2.63 | 313 | 0.956 | 0.072 | 0.046 |
| East | KY | 4.13 | 464 | 0.956 | 0.077 | 0.074 |
| East | MA | 5.55 | 549 | 0.952 | 0.075 | 0.071 |
| East | MUN | 1.00 | 107 | 0.954 | 0.046 | 0.000 |
| East | SC | 2.61 | 285 | 0.958 | 0.073 | 0.053 |
| East | UI | 5.67 | 621 | 0.957 | 0.076 | 0.089 |
| Chile | CH | 6.54 | 360 | 0.959 | 0.058 | 0.043 |

To generate sequence (haplotypic) datasets, we concatenated reads one and two and used the entire 295-bp sequences in these analyses. Loci were filtered to include only unlinked loci, but freely recombining within loci, represented in 90% of sampled individuals, resulting in a 165 locus dataset.

### Population genomic statistics

Summary statistics were generated using the *populations* script in STACKS and with the program Arlequin v3.5.2.2 (Excoffier and Lischer 2010). For each population we calculated observed homozygosity (H_obs_), nucleotide diversity (π), number of private loci, and the mean inbreeding coefficient per population (F_IS_). Pairwise F_ST_ values were calculated for all populations in Arlequin with 10,000 permutations for each to determine significance (P < 0.05). To visualize the F_ST_ matrix we created a UPGMA dendrogram and heatmap using the function *hclust* in the R package *ggdendro* (De Vries and Ripley 2016).

### Population structure

We used two different approaches to infer range-wide population structure and admixture of individuals. First, we used discriminant analysis of principal components (DAPC) in the R package *adegenet* (Jombart and Ahmed 2011) to partition genetic variation into clusters by maximizing variance between clusters and minimizing variance within clusters. We explored a range of cluster (*K*) numbers from 1 to 16 and used the estimated BIC from each model to identify the optimal *K*. To identify the optimal number of principal components retained in each analysis, we used cross-validation (xvalDapc with 10 retained). Scatterplots and barplots were used to visualize individual assignment to clusters.

Second, we used a model-based approach in the program ADMIXTURE v1.3 (Alexander et al., 2009), which uses a maximum-likelihood approach to assign individuals to genetic clusters. Five replicate analyses were performed for *K* = 1-16, with the optimal *K* chosen as having the lowest cross-validation error across replicates (Alexander and Lange 2011). We also evaluated levels of *K* above the identified optimum, as these may also be useful in explaining the data (Meirmans 2015). We used the web server CLUMPAK to generate assignment plots across multiple values of *K* (Kopelman et al., 2015).

### Analysis of Molecular Variance

Using the optimal clustering suggested by DAPC and Admixture analyses, we also performed an AMOVA with 1000 permutations, to estimate the degree of population structure and the covariance structure (1) within individuals in each sampled population, (2) within populations in estimated clusters, and (3) among clusters. Pairwise differentiation and its significance (computed as Weir and Cockerham’s Fst) between clusters was also estimated using 100 permutations.

### Contemporary gene flow estimation

To estimate contemporary gene flow between clusters identified by ADMIXTURE and DAPC, we performed a Bayesian Markov Chain Monte Carlo (MCMC) analysis using BayesAss3-SNPs (Mussmann et al., 2019), which implements an extension of BayesAss v.3.0 (Wilson and Rannala 2003) for large SNP datasets. We used 10 million iterations, discarding 1 million as burn-in, and sampled every 1000 iterations for estimates of posterior densities of migration estimates. Sufficient mixing of chains and convergence of MCMC were determined using TRACER v1.7.1 (Rambaut et al., 2018). Modes of posterior density estimates were obtained and compared across three runs (with different random number seeds).

### Ancestral Demographic model testing and gene flow estimation

Building on the resolution of genetic clusters, we used two approaches to infer population history and test models of gene flow across the range of *H. convergens*. First, we performed demographic model testing using a composite-likelihood modeling approach in fastsimcoal2 v2.5.2.21 (fsc2; Excoffier et al., 2013), which uses the site frequency spectrum (SFS) to infer demographic parameters. To construct the folded joint SFS for each population pair we used the *dadi.Spectrum.from_data_dict* command in dadi v1.6.3 (Gutenkunst et al., 2009). STACKS data filtering required a SNP to occur in ≥ 50% of individuals from each population, so the SFS was then down-projected to 50% to ensure the same number of gene copies were sampled for all loci. SFS entries with less than 10 SNPs were pooled (-*C10* option in fsc2). Invariable sites were not included in the SFS, so to scale estimation of other parameters from the SFS we fixed the ancestral population size to be 714,000 as calculated directly from our data (Excoffier et al., 2013; Lanier et al., 2015; Supporting information for details).

Population divergence models were constructed using the three population clusters identified in DAPC and Admixture analyses. A total of nine different models were generated covering various histories of population divergence, with and without migration, and with population sizes allowed to vary over time (Figure S1). In addition to these three-population models, we also tested pairwise population models, which reduced the number of estimated parameters and accounted for working with relatively small numbers of loci. Parameters estimated from models included the effective population size (N_e_) of each population, the divergence time for each splitting event (T_DIV_), and migration rates (m). We assumed a generation time of one year when converting estimates to years (Hagen 1962). Defined parameter ranges were uniformly distributed with N_e_ ranging from 10 to 10,000 and T_DIV_ from 10 to 10,000. A minimum of 100,000 simulations were performed to estimate the SFS, with a minimum and maximum of 10 and 100 loops (ECM cycles). The stopping criterion, defined as the minimum relative difference in parameters between two iterations, was set to 0.001. A total of 100 replicate runs were performed per model and the overall maximum likelihood (ML) run was retained for parameter estimation. The relative likelihood across compared models was generated using Akaike Information Criteria (AIC) as outlined in Excoffier et al., (2013). Confidence intervals for parameters in the best-supported model were obtained with a parametric bootstrap approach by simulating 100 SFS with the same number of SNPs from the ML point estimates.

*Second,* to estimate demographic parameters (effective population sizes, divergence times, and migration rates) under the most supported population tree from runs of *fastsimcoal2*, and to test scenarios of augmentation and gene flow between sampled populations of *H. convergens*, we utilized the parallel MCMC method implemented in IMa2p (Sethuraman et al., 2016, Hey and Nielsen 2007). IMa2p analyses use sequence data as input instead of SNP genotypes, so to perform these analyses we used a data set comprising 165 unlinked, and freely recombining-within loci (145 bases each). Assuming a population tree based on the sequence of branching events in the best-fitting model identified in fsc2, we performed a long run of the M mode (MCMC) with 150 chains distributed across 15 CPUs on the University of Kentucky’s Lipscomb High Performance Computing Cluster. The MCMC was allowed to burn-in for a period of 16 hours, and genealogies were sampled thereon for 32 hours after burn-in. Upper limits on prior values of all parameters were set based on the recommendations of Hey (2011) and adjusted (along with the length of runs) based on several short runs to ensure that the chains were mixing, swapping sufficiently, and reached stationarity prior to sampling. Marginal posterior density estimates of all parameters and 95% confidence intervals were computed on demographic scales by using the *Drosophila melanogaster* mutation rate of 3.5 x 10^-9^ per site per generation (Keightley et al., 2009). To test alternative models of gene flow, an additional L (likelihood ratio test) mode run was performed based on computing joint posterior densities by the method of Hey and Nielsen (2007) and Nielsen and Wakeley (2001). Three alternate demographic scenarios were tested against the fully nested model containing all bi-directional gene flow parameters between ancestral and contemporary populations: (1) a model of no immigration from eastern U.S. into western U.S. populations, (2) a model of no migration from Chile into the western U.S. populations, and (3) neither migration from Chile or Eastern populations into the Western populations.

## Results

### ddRAD dataset and summary statistics

We generated a total of ∼215 million paired sequence reads, with an average of 2,239,182 ± 1,244,709 reads per individual (Table S1). After excluding eight individuals with high missing data and filtering data as outlined above, we generated a data set comprising 9,824 bi-allelic SNPs (one SNP sampled per locus). After screening for loci with MAF < 0.05, and other filters, we also constructed a dataset with 1175 bi-allelic SNPs across a total of 78 individuals from 17 sampling locales. Generally, population genetic diversity as measured through summary statistics was comparable among all sampling sites (Table 1).

### Genetic structure and population differentiation

DAPC analysis assigned individuals to three genetic clusters (Figure 2) corresponding to (1) Arizona + California sites (hereafter referred to as the “western” cluster), (2) sites from the central and eastern US (east of the Rocky Mountains, hereafter referred to as the “eastern” cluster), and (3) a cluster comprising the Chilean site. The two commercially ordered populations from Arizona and California clustered with western populations. Individuals were assigned to their respective cluster with high probability. These geographic genetic clusters were genetically distinct from each other, with no overlap in ordination space.

**Figure 2:**
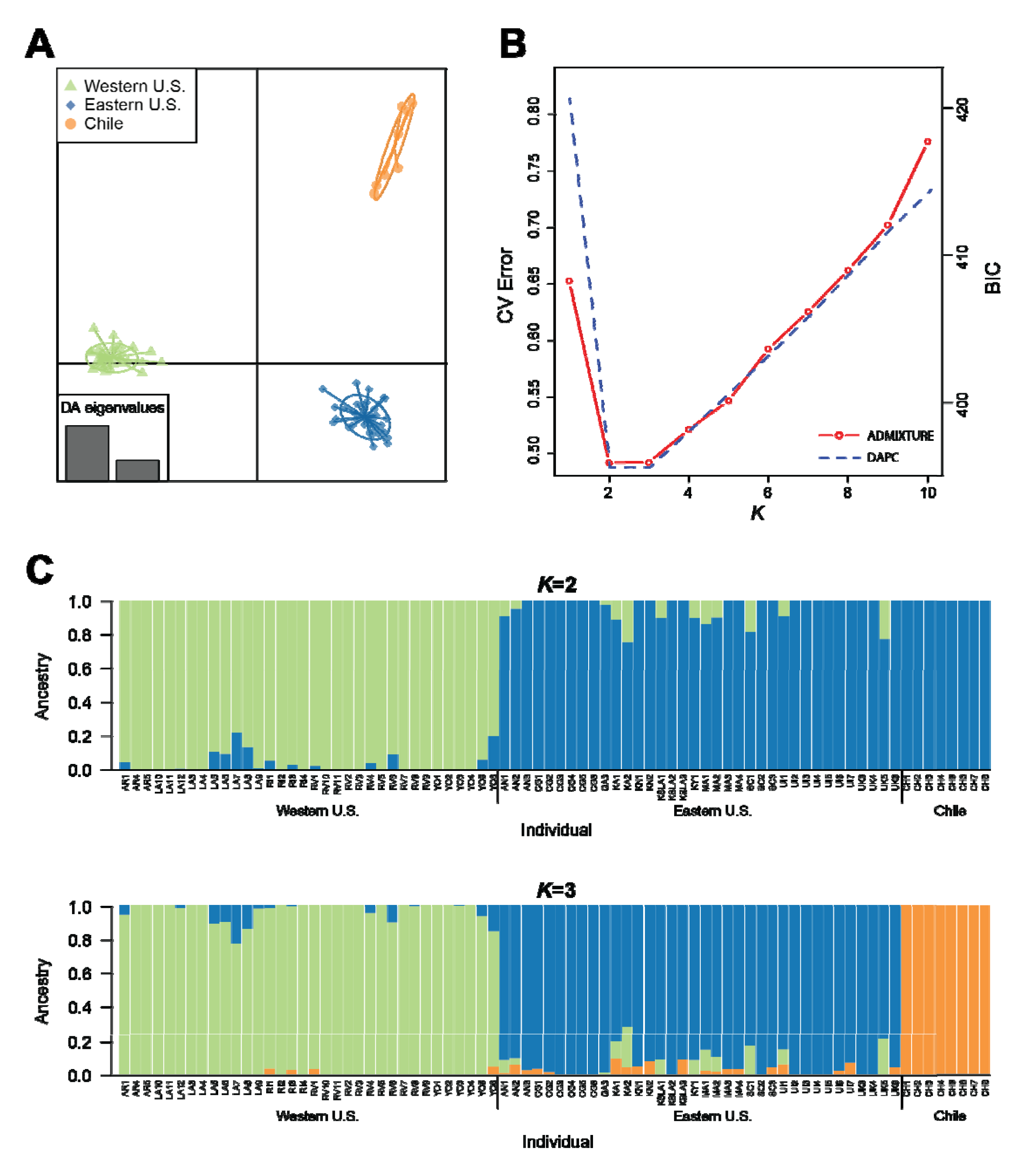
Population structure of *H. convergens*. (A) Scatter plot of *H. convergens* genotypes generated by DAPC at K = 3. (B) BIC values and cross validation errors for *K*=1-10, as estimated from DAPC and replicate runs of ADMIXTURE respectively. (C) Ancestry proportions plotted across all sequenced *H. convergens,* as estimated by ADMIXTURE (*K*=2 and 3)

ADMIXTURE analyses produced similar results (Figure 3). Cross-validation indicated *K* = 2 (c.v. error = 0.4920) as the optimal model with an eastern cluster and western + Chilean cluster. A *K* = 3 (c.v. error = 0.4922) model split the western and Chilean populations into separate clusters. Models of *K* ≥ 4 (c.v. error > 0.5214) produced new clusters that primarily created admixed individuals within the eastern and western populations with no further geographic partitioning, indicating that a *K* = 3 best represents the uppermost level of hierarchical geographic genetic structure across our study system.

**Figure 5:**
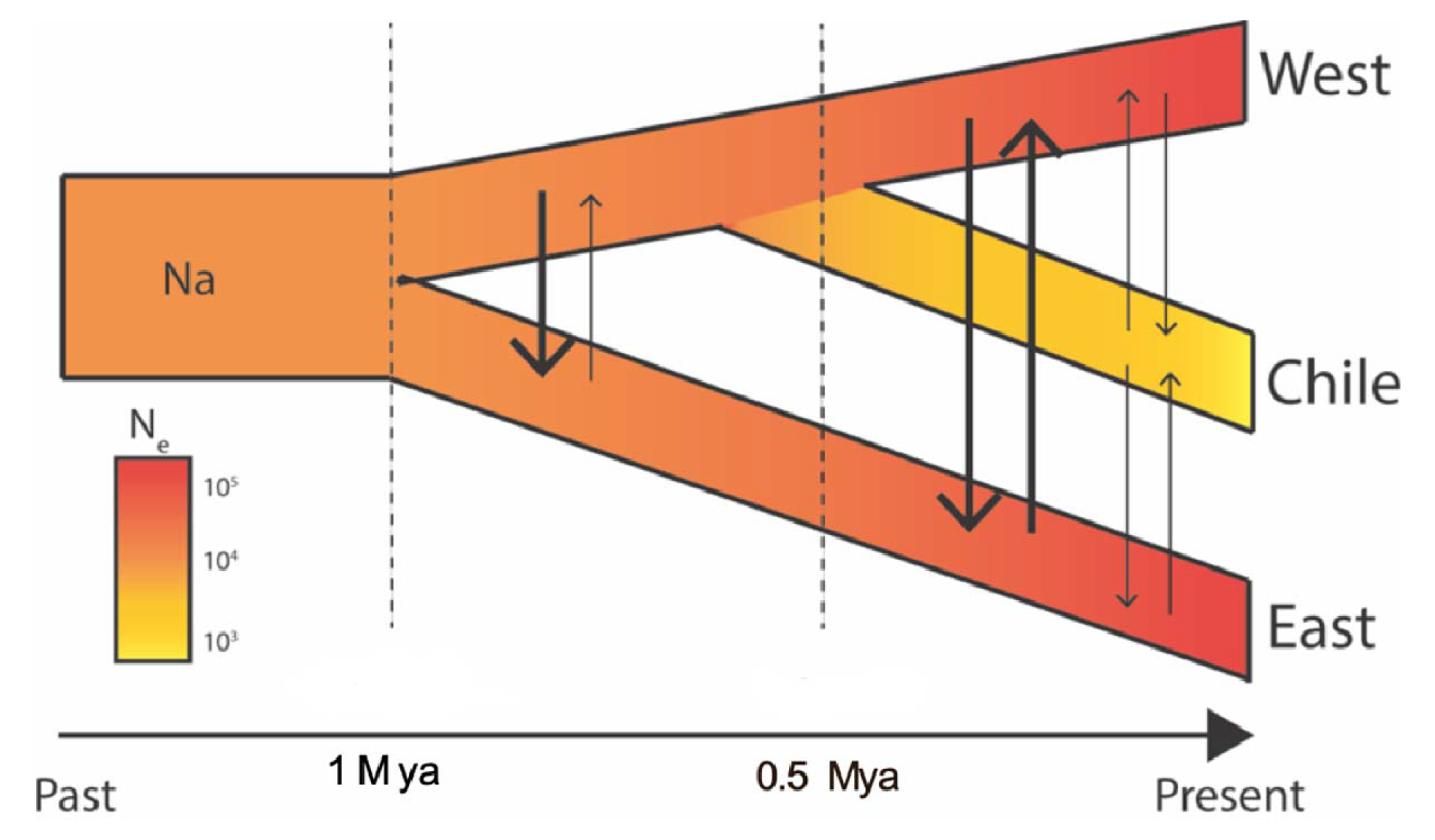
Best fitting topology and evolutionary model from the fsc2 analyses, showing an early split of the Eastern population of *H. convergens* from the common ancestor of the Western and Chilean populations, with some degree of continued asymmetric gene flow between them after divergence.

AMOVA analyses estimated that a majority of variation (59.96%, SS = 9255.21, df = 139) in the SNP data was explained by variability among individuals within all sampling sites. 38.51% of variation (SS = 4093.56, df = 2) was captured by variability among eastern, western, and Chilean clusters, while variation among populations within each cluster only contributed 1.53% of the total variation (SS = 1126.922, df = 14).

Pairwise F_ST_ values were not significantly different from zero for comparisons among sites within the eastern and western clusters (Figure S2; Table S3). However, F_ST_ values were significant for comparisons among sites between the three identified genetic clusters (F_ST_ = 0.13 - 0.49, p < 0.05), with the Chilean population identified as the most differentiated from Southern California (Riverside County).

### Contemporary gene flow

Estimates of contemporary gene flow using BayesAss3-SNPs indicated that the majority of gene flow occurred within identified clusters (Table 4), with a negligible fraction of migrants within each cluster classified as migrants. All three clusters had inbreeding coefficients of ≥ 0.15, indicating the presence of non-random mating within clusters, but little to no contemporary gene flow across clusters.

### Ancestral Demographic inference

Model testing in fsc2 favored a topology (model 2) with an early split between the eastern and western + Chile populations, followed by a subsequent split between the western and Chile population, and with the full set of migration parameters between both ancestral and contemporary populations (Fig. 4; Table 2). Model 2 was slightly favored over a full migration model that identified a more recent common ancestor between the eastern and western populations, with a more divergent Chilean population (Model 3; ΔAIC = 0.78). There was little support for a full migration model that identified a more recent common ancestor between the eastern and Chilean populations (Model 1; ΔAIC = 88.69). All evaluated models with a more restricted set of migration parameters (or no migration) received little support (ΔAIC > 141). This included models consistent with biocontrol efforts, in which gene flow moved unidirectionally from western populations into both eastern and Chilean populations.

**Table 2.** Relative likelihood of the assessed models of population history for *Hippodamia convergens*. Full descriptions of each model are provided in Figure S1.

| Model | Maximum likelihood | Number of Parameters (d) | AIC | ΔAIC | Model normalized relative likelihood |
| --- | --- | --- | --- | --- | --- |
| 1 | -7224.22 | 14 | 33296.76 | 88.69 | 3.29E-20 |
| **2** | **-7204.96** | **14** | **33208.08** | **0.00** | **5.96E-01** |
| 3 | -7205.13 | 14 | 33208.85 | 0.78 | 4.04E-01 |
| 4 | -7237.79 | 9 | 33349.25 | 141.17 | 1.32E-31 |
| 5 | -7240.98 | 9 | 33363.93 | 155.86 | 8.55E-35 |
| 6 | -7241.72 | 9 | 33367.34 | 159.26 | 1.56E-35 |
| 7 | -7340.25 | 7 | 33817.09 | 609.02 | 3.38E-133 |
| 8 | -7356.89 | 7 | 33893.74 | 685.66 | 7.68E-150 |
| 9 | -7306.14 | 7 | 33660.00 | 451.93 | 4.37E-99 |

Bayesian MCMC-based inference of demographic history under the most supported population tree from fastsimcoal2, using an isolation with migration model with IMa2p (Sethuraman and Hey 2015) estimated large bidirectional historical gene flow between eastern and western populations, and eastern and Chilean populations. No significant migration was estimated in either direction between the western and Chilean populations. Estimates of effective population sizes (population mutation rates) were concordant with fastsimcoal2, with the eastern and western populations estimated to be several orders larger than the Chilean population. Common ancestral populations were also estimated to be relatively small, indicative of a recent (post-divergence) population size increase in both eastern and western populations. In short, there is significant bidirectional historical migration between the eastern and western populations, and the eastern and Chilean populations (Table 5). Demographic estimates of effective population sizes, divergence times, and scaled migration rates were comparable between the results of fsc2, and IMa2p, with predominantly overlapping confidence intervals (Table 4), with relatively large eastern and western populations, compared to Chile.

**Table 3.** Demographic parameter estimates and confidence intervals (CIs) for *Hippodamia convergens* under the best-fitting model (model 2) from fastsimcoal2 (fsc2) and IMa2p. Parameter estimates in fsc2 were obtained as maximum likelihood estimates with CIs obtained by parametric bootstrapping. Parameter estimates in IMa2p were obtained as marginal posterior density estimates with CIs estimated as 95% highest posterior density intervals. Note that the Ancestral Ne was fixed in fsc2 analyses (denoted by *).

|  | **fsc2** | **IMA2p** |
| --- | --- | --- |
| # Loci (Type) | 4,139 (SNPs) | 165 (145 bp sequences) |
| $T_{DIV}$ East/West + Chile (years) | 1,037,392 (249,471-1,833,551) | 216,962 (177,514-256,510) |
| $T_{DIV}$ West/Chile (years) | 447,780 (82,360-593,672) | 78,895 (59,171-98,619) |
| $N_e$ Ancestral | 714,000* | 24,631 (14,778-54,187) |
| $N_e$ West+Chile | 184,264 (3,485-273,352) | 73,964 (39,447-108,481) |
| $N_e$ West | 2,524,867 (465,210-3,157,104) | 3,382,642 (2,593,688-3,372,781) |
| $N_e$ East | 1,653,579 (278,416-2,234,104) | 2,440,828 (2,115,384-2,948,718) |
| $N_e$ Chile | 76,408 (12,024-100,239) | 32,544 (24,654-42,406) |
| $N_m$ (E → Ancestor<sub>West + Chile</sub>) | 0.00 (0.00-0.08) | 0.135 (0.03-0.2) |
| $N_m$ (Ancestor<sub>West + Chile</sub> → E) | 0.00 (0.00-6.25) | 0.01 (0.00-4.41) |
| $N_m$ (W → E) | 6.80 (5.68-8.18) | 0.86 (0.47-2.67) |
| $N_m$ (E → W) | 0.00 (0.00-0.41) | 5.28 (3.96-7.75) |
| $N_m$ (W → C) | 0.00 (0.00-0.05) | 0.05 (0.01-0.08) |
| $N_m$ (C → W) | 0.85 (0.58-1.20) | 0.05 (0.01-1.09) |
| $N_m$ (E → C) | 0.25 (0.17-0.28) | 0.08 (0.06-0.11) |
| $N_m$ (C → E) | 0.83 (0.38-1.29) | 4.51 (2.58-6.11) |

**Table 4:**
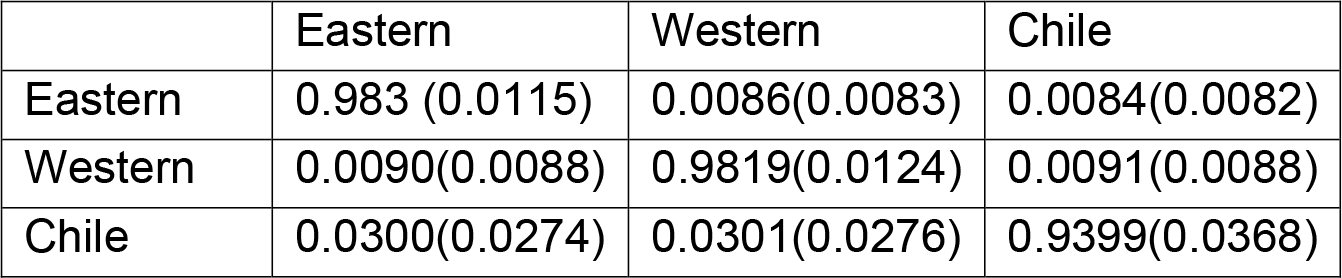
Mean contemporary migration rate estimates (fraction of individuals that are migrants from the cluster on the left to the cluster on the top), and standard deviations around the mean shown in parentheses between the three identified clusters of *Hippodamia convergens* (Eastern, Western, and Chilean). All estimates were performed using BA3-SNPS (Mussmann et al., 2019), which extends the method of BayesAss v.3.0 (Wilson and Rannala 2003).

**Table 5:** Likelihood ratio tests of different demographic models of *H. convergens* evolution, indicating support for "model 2" with significant bidirectional asymmetric historical migration rates.

| **H0 of Likelihood Ratio Test** | **-2LLR** | **df** | **p-value** | **Inference** |
| --- | --- | --- | --- | --- |
| No significant migration from East to West | 0.5874 | 1 | 0.44 | Inability to reject the H0 of no significant migration between Eastern and Western populations |
| No significant migration from Chile into the Western population | 131.4 | 1 | << 0.0001 | Reject the H0 of no significant migration from Chile into the Western population |
| No significant migration from Eastern to Western population, and no significant migration from Chile into the Western population | 276.1 | 2 | << 0.0001 | Reject the H0 of no significant migration into the Western population from either the Eastern or Chilean population |

## Discussion

The goal of this study was to investigate the impact of large-scale augmentative releases of *H. convergens* on its population structure and demographic history. Interestingly, our results indicate that releases with corresponding high gene flow from western into eastern populations have had little impact on population genetics of the species to date, with strong population structuring across the range and little signal of admixture of Western genotypes into Eastern populations. We discuss these findings in light of biological control and their implications for range wide population structure.

### Genetic Structure in US

Our results suggest historically strong genetic divergence between eastern U.S., western U.S., and Chilean populations of *H. convergens.* Estimates of contemporary gene flow indicated little to no continuing gene flow between our identified clusters. Meanwhile, demographic modeling of ancestral processes revealed population expansions of eastern and western populations with high gene flow from west into east, and a small, isolated Chilean population. We used two demographic inference approaches to explore ancestral population history (fastsimcoal2 and IMa2p), with comparable parameter estimates generated between models. However, uncertainty is associated with absolute values of parameter estimates from both programs which were scaled with a mutation rate from *Drosophila melanogaster*, and should be interpreted with caution. Instead, we focus on general patterns, which are corresponding between programs.

Both IMa2p and fastsimcoal2 analyses indicate asymmetrical gene flow between all population pairs. For fastsimcoal2 confidence intervals for migration were wide and often encompass zero, indicating low resolution of these parameters. We suspect that identification of migration may reflect low power to distinguish between models of isolation using a low number of SNPs, as simulation studies have found for IMa2 (Carstens et al., 2009; Hey et al., 2015). Population expansion over time is apparent in all analyses. These results are in contrast to an earlier microsatellite study, which detected admixture of Western genotypes into Eastern populations (Sethuraman et al., 2015). These results are especially interesting, considering that *Hippodamia convergens* is highly mobile, having been found to travel hundreds of miles (Hagen 1962). Formation of overwintering aggregations of adults occurs in late summer, with as many as 10^6^ beetles converging on a site (Baust and Morrissey 1975). Aggregations last from August to April, providing energy conservation for individuals with opportunities for mating (Szejner-Sigal & Williams 2022).

### Chilean Population Founding

We found the Chilean population has a much lower effective population size than eastern and western U.S. populations. This supports the hypothesis that the Chile population was recently introduced using commercially harvested populations from the Western USA (Hagen et al 1976). However, our estimates of recent (contemporary) gene flow using BayesAss indicates very little degree of continuing gene flow between the Chilean and mainland USA populations. These observations were also complemented by summary statistics (F_ST_) and population structuring programs (DAPC, ADMIXTURE), which detected strong population structure with little to no admixture between Eastern and Western populations. This suggests that commercially released Western collected individuals may not be successfully breeding and establishing gene flow in the East. Isolation of the Chilean population was evident, with many pairwise FST values > 0.40 (p < 0.05). This strong genetic population differentiation may be largely due to drift within the Chilean population, post-founding.

### Impact of Agriculture/Bio-control

Results from both IMa2p and fastsimcoal2 suggest that effective population size has increased over time across all structured populations of the species, and that the contemporary effective population sizes are large. This is contrasting to the previous microsatellite study finding population declines (Sethuraman et al., 2017), and speculation that native population of *H. convergens* have decreased after initialization of European agricultural processes, or in response to competition from invasive non-native predatory coccinellids, including the considerably larger Eurasian seven-spotted beetle, *Coccinella septempunctata*, and the Asian harlequin lady beetle, *Harmonia axyridis*. However, it is important to note that the large effective sizes that characterize both Eastern and Western US populations limit our ability to detect recent changes in population size. It is therefore possible that despite our results, populations could be currently in decline. The data collected here may therefore serve as a valuable baseline for genetic monitoring into the future (Schwartz et al., 2007).

### Importance for Biological Control

An appreciation of the evolutionary history of species that are manipulated by humans may be important for our understanding of the benefits, limitations and consequences of our activities. Following the large-scale changes in North American agroecosystems during the past 500 years, the native natural enemy fauna (e.g., arthropod predators) colonized these new agricultural systems to find prey. Thus, many species of North American insect predators have adapted to introduced agricultural crops from Europe and Asia, but our understanding of the genetic bases of these adaptations is limited.

Augmentative releases of *H. convergens* represents a unique example of biological control, and allows for the examination of several potential ecological and genetic non-target effects of these releases on other predatory species and local populations of *H. convergens* (O’Neil et al 1998; Saito and Bjornson 2008, 2013). Many augmentative releases of *H. convergens* have appropriately targeted pest species in agricultural systems in California and Arizona (Hagen 1962; Dreistadt and Flint 1996; Flint and Dreistadt 2005; Hagler 2009). But, for at least the past 75 years, field collected *H. convergens* from the western United States have been sold and released east of the Rocky Mountains.

Our analysis of SNP variation of North American *Hippodamia convergens* indicates that historical events (e.g. glaciation, introduction of western European agriculture) and not recent human movements of western USA populations of *H. convergens* explain the current distribution and population structure of this predatory insect species. Thus, the current distribution and population structure of *H. convergens* in North America can be explained from historical events, our analyses showed that annual augmentative releases are not affecting local populations. Our previous studies documented plasticity in the response of Eastern and Western populations of *H. convergens* to aphid prey with no hybrid vigor or heterozygote advantages in hybrids (Grenier et al 2021). Based on our results, we tentatively propose that augmentative releases of western US populations of *H. convergens* do not have adverse non-target genetic effects on eastern US populations. However, the potential long-term ecological non-target effects of distributing pathogens and parasitoids within adult *H. convergens* from western collection sites on eastern populations of ladybeetle populations remains to be determined (Bjornson 2008, Sato & Bjornson 2008).

### Data Availability

All sequences generated by this study have been deposited in SRA (accession ID:), and the final SNP data can be accessed via Datadryad (link). All scripts and code used in population genomic analyses have been deposited on GitHub (www.github.com/arunsethuraman/hconddrad).

## Acknowledgments

This work was supported by NSF ABI 1564659, NSF CAREER 2042516 to AS. This work was funded by the National Institute of Food and Agriculture, U.S. Department of Agriculture, Hatch Program under accession number 1008480 and funds from the University of Kentucky Bobby C. Pass Research Professorship to JJO. This research was supported in part by a Research Support Grant from the University of Kentucky Office of the Vice President for Research to DWW and JJO. This research includes calculations carried out on HPC resources supported in part by the National Science Foundation through major research instrumentation grant number 1625061 and by the US Army Research Laboratory under contract number W911NF- 16-2- 0189.

---

**Figure S1:**
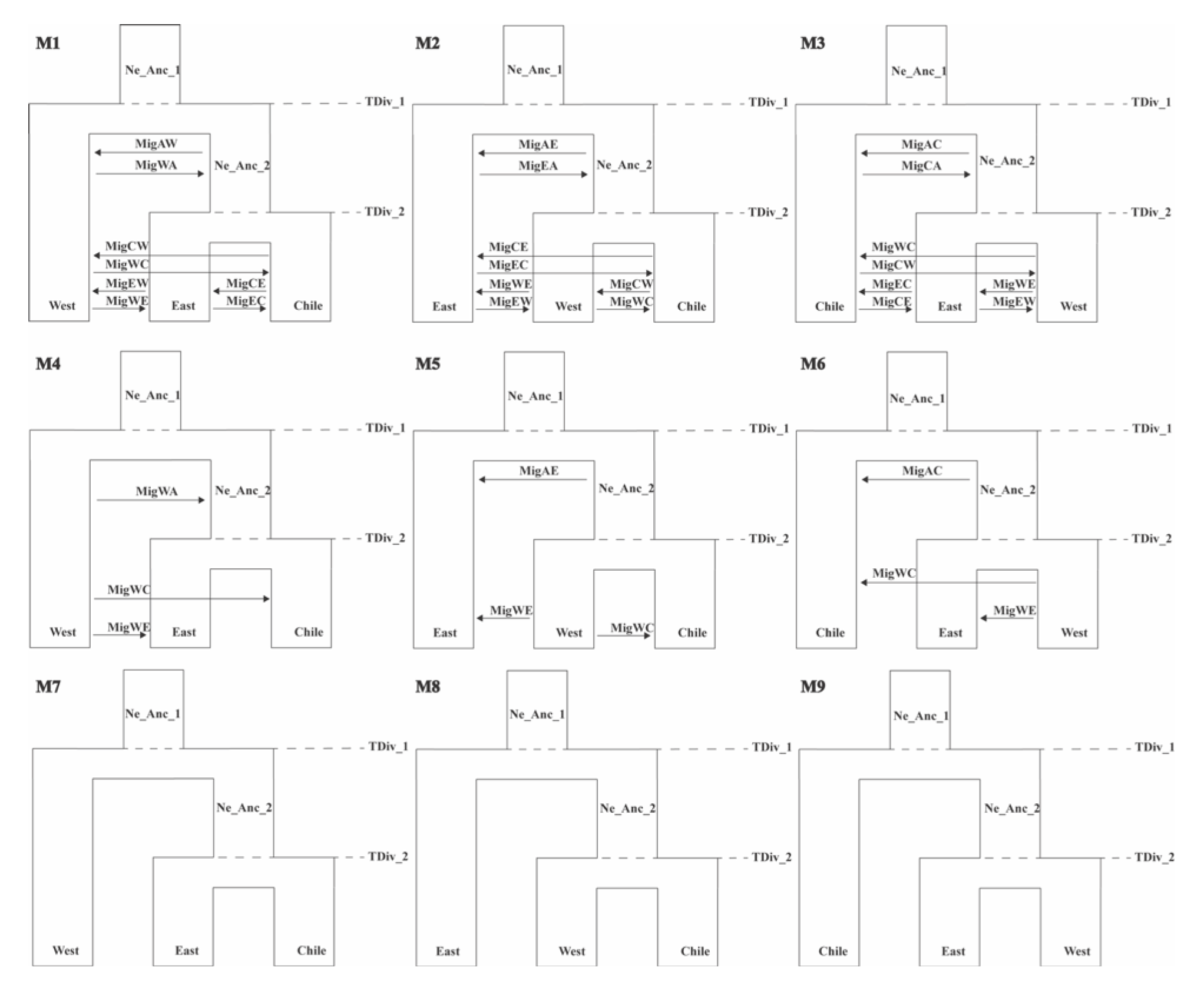
All evolutionary models and competing topologies tested in FSC2 under the three population model as estimated by ADMIXTURE.

**Figure S2:**
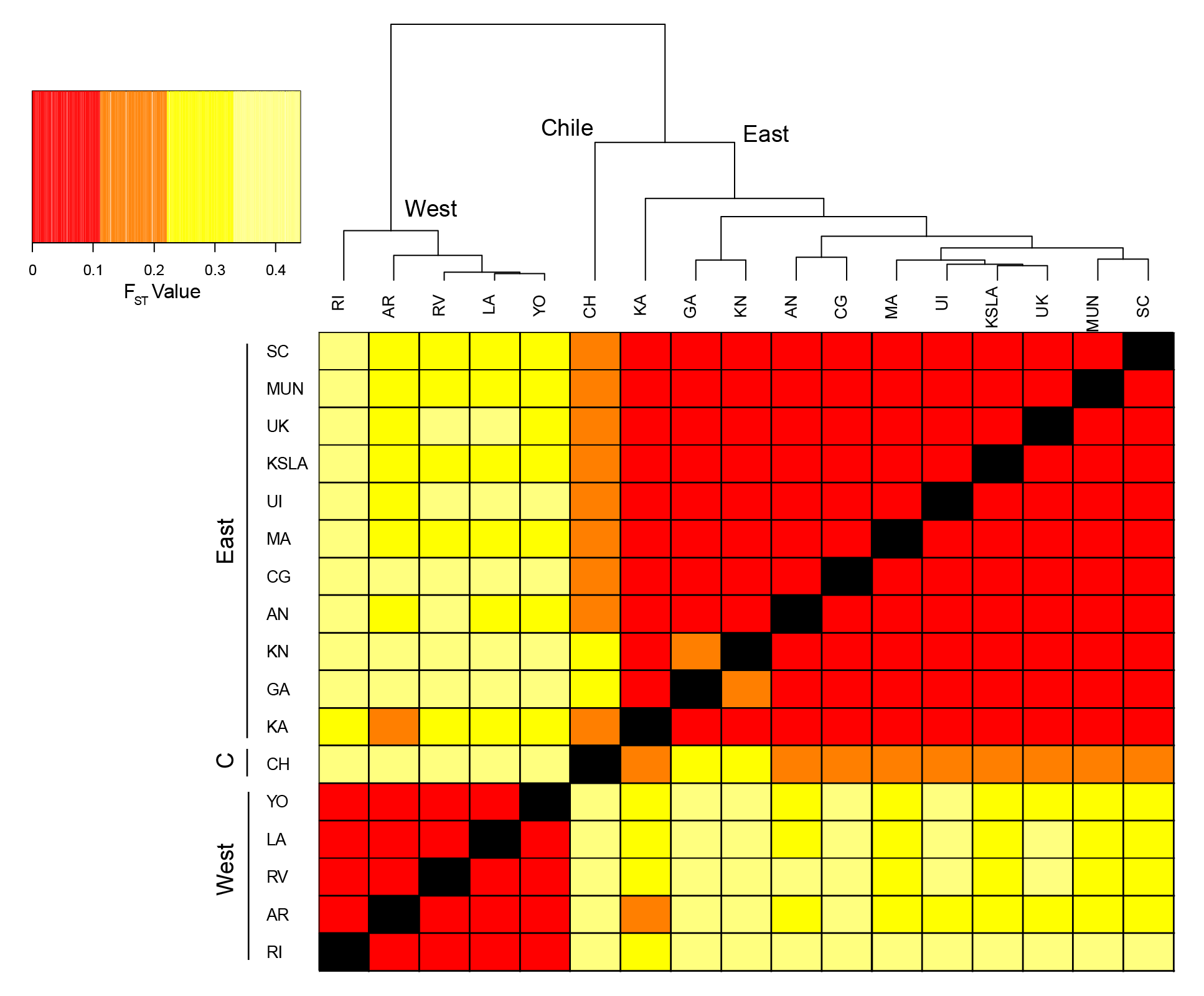
Estimates of pairwise Weir and Cockerham’s Fst across all sampled *H. convergens* populations.

**Figure S3:**
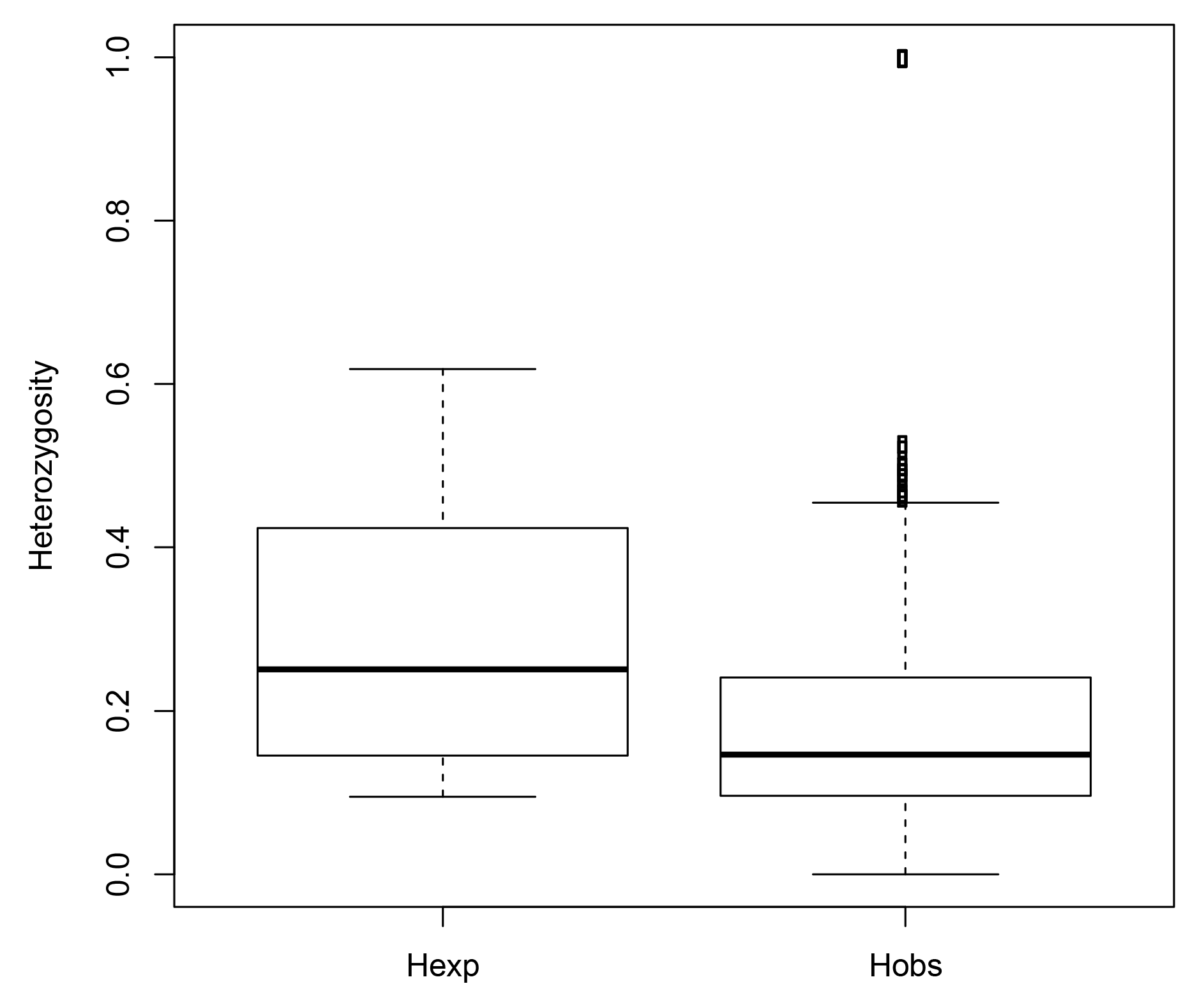
Expected and observed heterozygosities across all loci, all populations of *H. convergens* used in this study.

**Figure S4:**
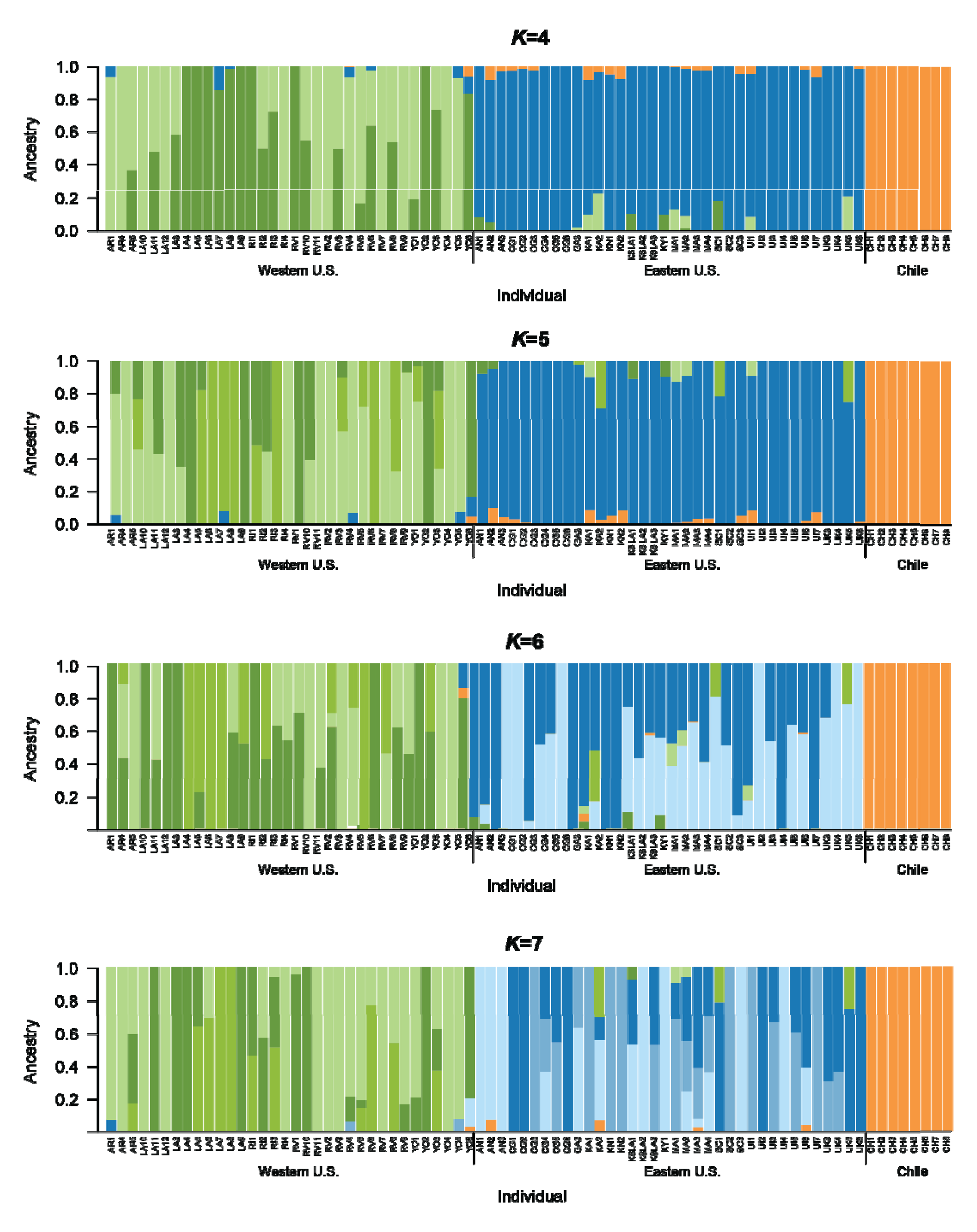
Ancestry proportions plotted across all sequenced *H. convergens*, as estimated by ADMIXTURE (K=4,5,6,7)

**Table S1.** Sampling localities, GPS Coordinates, and collection information for *H. convergens* individuals.

| Individual Symbol | Location | Lat | Long | Date Collected | # of Samples Sequenced | # of Samples Dropped |
| --- | --- | --- | --- | --- | --- | --- |
| AN | Ankeny, Iowa | 41.74°N | 93.6°W | May 1992 | 3 |  |
| CG | Coffee County, GA | 31.56°N | 82.84°W | Aug 2011 | 6 |  |
| GA | Tifton, GA | 31.45°N | 83.52°W | Apr 1994 | 1 |  |
| KA | Kadoka, SD | 43.84°N | 101.51°W | Jun 1993 | 2 |  |
| KN | Knoxville, Iowa | 41.32°N | 93.11°W | May 1992 | 2 |  |
| KSLA | Laurence, Kansas | 36.97 N | 95.23 W | May 2015 | 3 |  |
| KY* | UK North Farm | 38.14 N | 84.51 W | May 2015 | 1 |  |
| MA | Manhattan, Kansas | 39.19°N | 96.57°W | May 2011 | 11 | 7 |
| MC | Mitchell County, Iowa | 43.36 N | 92.77 W | ?? | 1 | 0 |
| MUN | Muncie, Indiana | 40.19 N | 85.40 W | ?? | 1 |  |
| SC | Tall Grass Prairie Site, Strong City Kansas | 38.39 N | 96.53 W | May 2015 | 3 |  |
| UI | Urbandale, Iowa | 41.63°N | 93.71°W | Oct 2015 | 7 |  |
| UK* | UK North Farm, KY | 38.14°N | 84.51°W | Oct 2011 | 4 |  |
| AR | Arbico Organic in Tucson, AZ | 32.45°N | 110.79°W | May 2011 | 6 | 3 |
| LA | Angeles National Forest, Los Angeles City, CA | 34.44°N | 118.11°W | Oct 2011 | 10 |  |
| RI | San Jacinto Mountain Reserve, Riverside City, CA | 33.81°N | 116.68°W | Oct 2011 | 4 |  |
| RV | Rincon-Vitova Insectary in Ventura, CA | 34.29°N | 119.29°W | Oct 2015 | 12 |  |
| YO | Yolo County, CA | 38.7°N | 121.87°W | Apr 2012 | 6 |  |
| CH | Santiago, Chile | 33.45°S | 70.67°W | Nov 2011 | 8 |  |

\*Samples pooled for all analyses

**Table S2.** Illumina sequence reads for each individual.

| Individual | Total | No | Low | Reads |
| --- | --- | --- | --- | --- |

|  | Reads | RadTag | Quality | Retained |
| --- | --- | --- | --- | --- |
| 1c | 2995968 | 15026 | 14385 | 2966557 |
| 1cA | 4900528 | 11214 | 26659 | 4862655 |
| **2c** | 409710 | 8143 | 2352 | 399215 |
| 3c | 1322626 | 7005 | 6232 | 1309389 |
| 3cA | 3990640 | 9316 | 18430 | 3962894 |
| 4c | 838298 | 7569 | 3830 | 826899 |
| 4cA | 1481192 | 7577 | 6916 | 1466699 |
| 5c | 2796856 | 12546 | 13442 | 2770868 |
| 5cA | 4924756 | 14518 | 24399 | 4885839 |
| AN1 | 3391414 | 5936 | 15768 | 3369710 |
| AN2 | 2980832 | 5104 | 14058 | 2961670 |
| AN3 | 3166168 | 17209 | 14974 | 3133985 |
| **AR1** | 827064 | 5875 | 4061 | 817128 |
| **AR2** | 485126 | 4156 | 2688 | 478282 |
| **AR3** | 271230 | 3749 | 1503 | 265978 |
| AR4 | 1155104 | 4703 | 6518 | 1143883 |
| AR5 | 2002780 | 8913 | 11761 | 1982106 |
| CG1 | 2447188 | 5931 | 12285 | 2428972 |
| CG2 | 2919260 | 7477 | 15023 | 2896760 |
| CG3 | 4336964 | 18026 | 21855 | 4297083 |
| CG4 | 4324316 | 12041 | 20917 | 4291358 |
| CG5 | 3968298 | 11291 | 17045 | 3939962 |
| CG6 | 2944178 | 9473 | 13610 | 2921095 |
| CH1 | 4169932 | 12664 | 20263 | 4137005 |
| CH2 | 5284496 | 9539 | 24405 | 5250552 |
| CH3 | 1918404 | 19203 | 10024 | 1889177 |
| CH4 | 2191154 | 9547 | 10450 | 2171157 |
| CH5 | 2332586 | 8261 | 11598 | 2312727 |
| CH6 | 1169270 | 16902 | 5227 | 1147141 |
| CH7 | 760818 | 9156 | 3747 | 747915 |
| CH8 | 690512 | 4839 | 4269 | 681404 |
| GA3 | 2334332 | 10472 | 11587 | 2312273 |
| KA1 | 1494258 | 5525 | 7374 | 1481359 |
| KA2 | 1636128 | 4215 | 8029 | 1623884 |
| KN1 | 2000726 | 19172 | 10094 | 1971460 |
| KN2 | 1595216 | 5770 | 8670 | 1580776 |
| KSLA1 | 4536300 | 25244 | 21632 | 4489424 |
| KSLA2 | 4191376 | 10986 | 19410 | 4160980 |
| KSLA3 | 4882862 | 9416 | 24239 | 4849207 |
| KY1 | 1911680 | 8857 | 10110 | 1892713 |
| LA10 | 1088236 | 8631 | 6063 | 1073542 |
| LA11 | 1193658 | 4599 | 5469 | 1183590 |
| LA12 | 1323050 | 18837 | 6262 | 1297951 |
| LA3 | 2055932 | 10258 | 9592 | 2036082 |
| LA4 | 1564552 | 14906 | 7885 | 1541761 |
| LA5 | 2604182 | 6063 | 12019 | 2586100 |
| LA6 | 3126562 | 13735 | 15818 | 3097009 |
| LA7 | 3596796 | 7825 | 17569 | 3571402 |
| LA8 | 1986974 | 6832 | 9791 | 1970351 |
| LA9 | 1251152 | 6332 | 6731 | 1238089 |
| MA1 | 754216 | 6956 | 4057 | 743203 |
| MA10 | 2665060 | 9051 | 12306 | 2643703 |
| MA2 | 1078174 | 5235 | 5374 | 1067565 |
| **MA3** | 616860 | 4890 | 2600 | 609370 |
| **MA4** | 514322 | 3917 | 3038 | 507367 |
| **MA5** | 436012 | 3987 | 3283 | 428742 |
| **MA6** | 215626 | 5578 | 1372 | 208676 |
| MA7 | 2299890 | 5754 | 11342 | 2282794 |
| MA8 | 2248204 | 14240 | 10771 | 2223193 |
| MA9 | 2460212 | 9111 | 12060 | 2439041 |
| RI1 | 2526660 | 9284 | 14300 | 2503076 |
| RI2 | 2153790 | 6611 | 9397 | 2137782 |
| RI3 | 3107266 | 20771 | 15628 | 3070867 |
| RI4 | 3564984 | 10710 | 17684 | 3536590 |
| RV1 | 2308182 | 7433 | 11350 | 2289399 |
| RV10 | 3714426 | 6674 | 16372 | 3691380 |
| RV11 | 2384022 | 7732 | 11612 | 2364678 |
| RV2 | 2999400 | 7492 | 14890 | 2977018 |
| RV3 | 1954652 | 5074 | 10554 | 1939024 |
| RV4 | 1674432 | 11605 | 8097 | 1654730 |
| RV5 | 3310764 | 9219 | 15437 | 3286108 |
| RV6 | 1393418 | 5985 | 6769 | 1380664 |
| RV7 | 2308926 | 7044 | 11112 | 2290770 |
| RV8 | 3286036 | 6608 | 16453 | 3262975 |
| RV9 | 3991772 | 6476 | 20465 | 3964831 |
| SC1 | 2277802 | 19421 | 10590 | 2247791 |
| SC2 | 4340706 | 10942 | 24803 | 4304961 |
| SC3 | 4713502 | 7602 | 23854 | 4682046 |
| UI1 | 2015532 | 7586 | 9519 | 1998427 |
| UI2 | 1585724 | 19167 | 7867 | 1558690 |
| UI3 | 1542580 | 5621 | 7628 | 1529331 |
| UI4 | 1378732 | 5529 | 6764 | 1366439 |
| UI5 | 1001040 | 5139 | 4811 | 991090 |
| UI6 | 1357690 | 6206 | 6523 | 1344961 |
| UI7 | 1449640 | 6553 | 6875 | 1436212 |
| UK3 | 1230242 | 30961 | 6232 | 1193049 |
| UK4 | 2891826 | 6663 | 13152 | 2872011 |
| UK5 | 1981814 | 6934 | 9538 | 1965342 |
| UK6 | 1951562 | 8059 | 9314 | 1934189 |
| YO1 | 1406014 | 13654 | 7250 | 1385110 |
| YO2 | 1644546 | 13515 | 7606 | 1623425 |
| YO3 | 1016842 | 6998 | 5101 | 1004743 |
| YO4 | 1161226 | 18426 | 5070 | 1137730 |
| YO5 | 1635688 | 7667 | 7587 | 1620434 |
| YO6 | 2554342 | **7977** | 12472 | 2533893 |

Individuals in bold excluded from analysis due to excessive missing genotypes

**Table S3.**
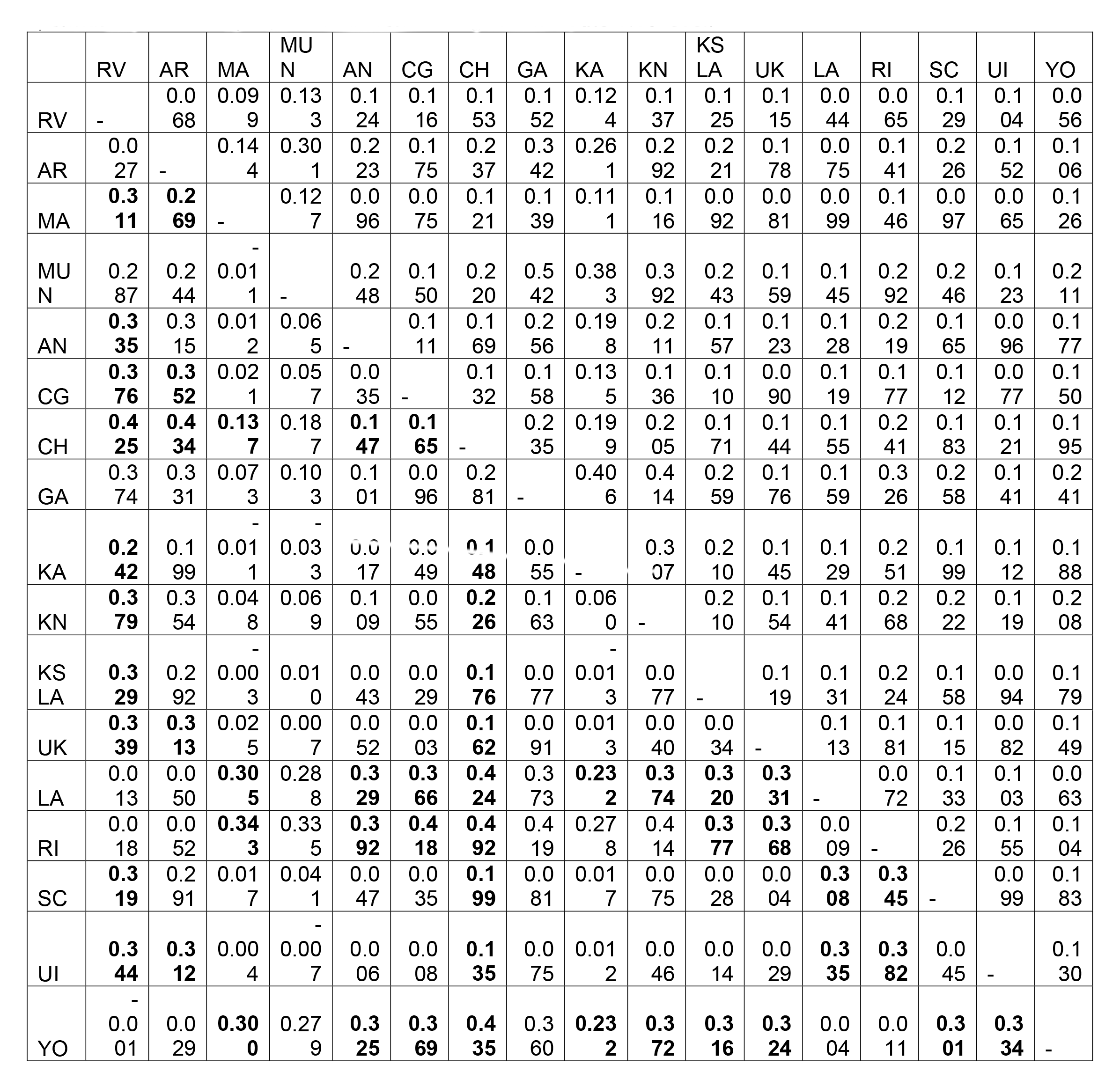
Pairwise F_ST_ values as calculated by STACKS (above the diagonal) and Arlequin (below the diagonal) for 17 populations of *H. convergens*. Absolute values differ because of differences in how the programs treat missing data and nonpolymorphic sites. Significance was determined with 10,000 permutations in Arlequin, values in bold are significant at the 0.05 level.

